# Evolution of Nuclear Receptors in Platyhelminths

**DOI:** 10.1101/2021.04.14.439782

**Authors:** Wenjie Wu, Philip T. LoVerde

## Abstract

Since the first complete set of Platyhelminth nuclear receptors (NRs) from *Schistosoma mansoni* were identified a decade ago, more flatworm genome data is available to identify their NR complement and to analyze the evolutionary relationship of Platyhelminth NRs. NRs are important transcriptional modulators that regulate development, differentiation and reproduction of animals. In this study, NRs are identified in genome databases of thirty-three species including in all Platyhelminth classes (Rhabditophora, Monogenea, Cestoda and Trematoda). Phylogenetic analysis shows that NRs in Platyhelminths follow two different evolutionary lineages: 1) NRs in a free-living freshwater flatworm (*Schmidtea mediterranea*) and all parasitic flatworms share the same evolutionary lineage with extensive gene loss. 2) NRs in a free-living intertidal zone flatworm (*Macrostomum lignano*) follow a different evolutionary lineage with a feature of multiple gene duplication and gene divergence. The DNA binding domain (DBD) is the most conserved region in NRs which contains two C4-type zinc finger motifs. A novel zinc finger motif is identified in parasitic flatworm NRs: the second zinc finger of parasitic Platyhelminth HR96b possesses a CHC2 motif which is not found in NRs of all other animals. In this study, novel NRs (members of NR subfamily 3 and 6) are identified in flatworms, this result demonstrates that members of all six classical NR subfamilies are present in the Platyhelminth phylum. NR gene duplication, loss and divergence in Platyhelminths are analyzed along with the evolutionary relationship of Platyhelminth NRs.

## INTRODUCTION

Platyhelminths (flatworms) are one of the largest animal phyla, it includes more than 20,000 species [1, 2]. Some flatworms are important disease-causing parasites of humans and livestock, e.g. Schistosomiasis, Paragonimiasis and Cestodiasis. Platyhelminths are bilaterally symmetrical, non-segmented acoelomates without an anus. Although they have an excretory system they lack respiratory and circulatory systems, all flatworms are hermaphroditic except Schistosoma sp. where asexual and sexual reproduction are present [3]. Platyhelminths are traditionally divided into four classes: Rhabditophora, Monogenea, Cestoda (tapeworms) and Trematoda (flukes). The class Rhabditophora includes all free-living flatworms, while all members in classes of Monogenea, Trematoda, and Cestoda are parasitic flatworms. From an evolutionary point of view, the parasitic classes arose from a primitive free-living flatworm [3].

Nuclear receptors (NRs) are important transcriptional modulators that regulate development, differentiation and reproduction of animals. Most of NRs share a common tertiary structure: A/B-C-D-E domains. The A/B domain is highly variable, the C domain is the DNA-binding domain (DBD) which is the most conserved region containing two zinc finger motifs, the D domain a flexible hinge between the C and E domains and is poorly conserved, the E domain contains the ligand binding domain (LBD) that is involved in transcriptional activation. Atypical NRs are found in some animals, e.g. NRs with a DBD but no LBD are found in arthropods and nematodes, members without a DBD but with a LBD are present in some vertebrates, and NRs with two DBDs and a single LBD (2DBD-NRs) are identified in protostomes. A phylogenetic analysis of the NRs divides them into six classical subfamilies by alignment of the conserved DBD [4]. The early study of the complete genome sequence of ecdysozoan Arthropods (*Drosophila melanogaster*) [5] the mosquito (*Anopheles gambiae*) [6], free-living nematodes (*Caenorhabditis elegans* and *C. briggsae*) [7, 8], tunicata (*Ciona intestinalis*) [9], mammalians: rat (*Rattus norvegicus*), mouse (*Mus musculus*) [10] and human (*Homo sapiens*) [11]) revealed that NRs in insects, tunicata and mammalians share the same evolutionary lineage with extensive gene loss/duplication, while NRs in nematodes follow a different evolutionary lineage with a feature of multiple duplication of SupNRs and gene loss.

Since we identified the first complete set of Platyhelminths nuclear receptors (NRs) from *Schistosoma mansoni* a decade ago [12], more flatworm genome data has become available to identify their NR complement. Analysis of the NR complement of the blood fluke *Schistosoma mansoni* [12-14] and tapeworm *Echinococcus multilocularis* [13-15] shows that NRs in *S. mansoni* and *E. multilocularis* share the same evolutionary lineage as that of Deuterostomia and the arthropods in the Ecdysozoan clade of the Protostomia, but some divergent NRs in flatworms are not present in Deuterostomia or/and arthropods, for example 2DBD-NRs [16]. In this study, we analyzed genome databases of Platyhelminth Wormbase website including species from all of the four classes in Platyhelminths [17, 18]. Identification of the NR complement will contribute to a better understanding of the evolution of NRs in Platyhelminths.

## MATERIALS AND METHODS

### 1. Data mining

Nuclear receptors in Platyhelminths were mined from the genome databases in Wormbase website [17, 18]. Amino acid sequence of DBD of SmTRa, SmHR96b and Sm2DBDα were used as the query to tbalstn against all available databases. Any sequence that contains a zinc finger structure of the NR DBD (Cys-X2-Cys-X13-Cys-X2-Cys or Cys-X5-Cys-X9-Cys-X2-Cys) was retained, and the deduced amino acid sequence of the DBD was obtained by the conserved junction position (JP) and GT-AG rule if there was an intron in the DBD coding region. The analyzed Platyhelminth genome databases are listed in **Table 1**.

**Table 1.** Nuclear receptors analyzed in Platyhelminth species.

| CLASS | ORDER | FAMILY | SPECIES<br>REFERENCES | SHOR DESCRIPTION |
| --- | --- | --- | --- | --- |
| Rhabditophora | Rhabditophora | Rhabditophorae | *Macrostomum lignano* (Ml) [22, 23] | Free-living, intertidal flatworm that can regenerate most of its body parts. |
|  | Tricladida | Dugesidae | *Schmidtea mediterranea* (Sme) [24-26] | Free-living, freshwater flatworm that can regenerate an entire organism. |
| Monogenea | Monopisthocotylea | Gyrodactylidae | *Gyrodactylus salaris* (Gs) [27] | Salmon fluke that lives on the body surface of freshwater fish. |
|  | Polyopisthocotylea | Polystomatidae | *Protopolystoma xenopodis* (Px) [28] | Fluke of African clawed frogs where the adult worms live in the host's urinary bladder. |
| Cestoda | Cyclophyllidea | Hymenolepididae | *Hymenolepis diminuta* (Hd) [28] | Rat tapeworm. The intermediate hosts: arthropod; the definitive hosts: rodent. |
|  |  |  | *H. microstoma* (Hm) [29] | Rodent tapeworm. The intermediate hosts: arthropod; the definitive hosts: rodent. |
|  |  |  | *H. nana* (Hn) [28] | Rodent tapeworm. The intermediate hosts: arthropod; the definitive hosts: human and rodent. |
|  |  | Taeniidae | *Echinococcus Canadensis* (Ec) [30] | Canid tapeworm. The intermediate hosts: cervid, camels, cattle etc.; the definitive host: dogs and other canids. |
|  |  |  | *E. granulosus* (Eg) [29, 31] | Dog tapeworm. The intermediate hosts: human, horse; the definitive host: dogs. |
|  |  |  | *E. multilocularis* (Em) [29] | Fox tapeworm. The intermediate hosts: wild rodents; the definitive hosts: foxes, coyotes, dogs and other canids. |
|  |  |  | *Taenia asiatica* (Ta) [32] | Human tapeworm. The intermediate hosts: pigs; the definitive host: human. |
|  |  |  | *T. multiceps* (Tm) [33] | Dog tapeworm. The intermediate hosts: sheep, goat and cattle; the definitive hosts: Canids. |
|  |  |  | *T. saginata* (Ts) [32] | Beef tapeworm. The intermediate hosts: Cattle; the definitive hosts: human. |
|  |  |  | *T. solium* (Tso) [29] | Pork tapeworm. The intermediate host: pigs; the definitive host: human. |
|  |  |  | *Hydatigera taeniaeformis* (Ht) [28] | Cat tapeworm. The intermediate hosts: rodents; the definitive hosts: cats. |
|  |  | Mesocetoididae | *Mesocetoides corti* (Mc) [28] | Mouse tapeworm. The intermediate host: birds and small mammals; the definitive host: cats and dogs. |
|  | Diphyllobotridea | Diphyllobothriidae | *Diphyllobothrium latum* (Dl) [28] | Human tapeworm. The intermediate hosts: fish; the definitive hosts: human. |
|  |  |  | *Schistocephalus solidus* (Ss) [28] | Bird tapeworm. The first intermediate hosts: cyclopoid copepod; the second intermediate hosts: fish; the definitive hosts: fish-eating water bird. |
|  |  |  | *Spirometra erinaceieuropaei* (Se) [34] | Tapeworm. The first intermediate hosts: cyclopoid copepod; the second intermediate hosts: fish, reptiles, or other amphibians; the definitive hosts: cats and dogs. |
| Trematoda | Opisthorchida | Opisthorchiidae | *Clonorchis sinensis* (Cs) [35] | Human liver fluke. The first intermediate hosts: freshwater snails; the second intermediate hosts: freshwater fish; the definitive hosts: human and other mammals. |
|  |  |  | *Opisthorchis felineus* (Of) [36] | Cat liver fluke. The first intermediate hosts: freshwater snails; the second intermediate hosts: freshwater fish; the definitive hosts: human and other mammals. |
|  |  |  | *O. viverrini* (Ov) [37] | Human liver fluke. The first intermediate hosts: freshwater snails; the second intermediate hosts: freshwater fish; the definitive hosts: human and other mammals. |
|  | Echinostomida | Echinostomatidae | *Echinostoma caproni* (Eca) [28] | Intestinal fluke. The first intermediate hosts: freshwater snails; the second intermediate hosts: freshwater snails or frogs. Humans can be the definitive host. |
|  |  | Fasciolidae | *Fasciola hepatica* (Fh) [38, 39] | Liver fluke. The intermediate hosts: freshwater snails; The definitive hosts: cattle, sheep and other mammals including human. |
|  | Strigeidida | Schistosomatidae | *Schistosoma bovis* (Sb) [40] | Blood fluke. The intermediate hosts: freshwater snails; the definitive host: ruminants (cattle, goats, sheep, etc). |
|  |  |  | *S. curassoni* (Sc) [28] | Blood fluke. The intermediate hosts: freshwater snails; the definitive hosts: ruminants (cattle, goat, sheep, etc). |
|  |  |  | *S. haematobium* (Sh) [41] | Blood fluke. The intermediate hosts: freshwater snails; the definitive hosts: cattle, goat or sheep. |
|  |  |  | *S. japonicum* (Sj) [42] | HZoonotic blood fluke. The intermediate hosts: freshwater snails; the definitive hosts: human. |
|  |  |  | *S. mansoni* (Sm) [43] | Human blood fluke. The intermediate hosts: freshwater snails; the definitive hosts: human. |
|  |  |  | *S. margrebowiei* (Sma) [28] | Blood fluke. The intermediate hosts: freshwater snails; the definitive hosts: antelope, buffalo and waterbuck. |
|  |  |  | *S. matthei* (Sma) [28] | Blood fluke. The intermediate hosts: freshwater snails; the definitive hosts: bovid. |
|  |  |  | *S. rodhaini* (Sr) [28] | Blood fluke. The intermediate hosts: freshwater snails; the definitive host: rodent. |
|  |  |  | *Trichobilharzia regenti* (Tr) [28] | Neuropathogenic fluke. The intermediate host: freshwater snail; the definitive host: avian. |

### 2. Phylogenetic analysis

Phylogenetic trees were constructed from deduced amino acid sequences of the DBD and/or ligand binding domain (LBD), the sequences are aligned with ClustalW [19], phylogenetic analysis of the data set is carried out using Bayesian inference (MrBAYES v3.1.1) [20] with a mix amino acid replacement model + gamma rates. The trees were started randomly; four simultaneous Markov chains were run for 5 million generations, the trees were sampled every 100 generations, the Bayesian posterior probabilities (PPs) were calculated using a Markov chain Monte Carlo (MCMC) sampling approach implemented in MrBAYES v3.1.1, with a burn-in value setting at 12,500. The same data set was also tested by Maximum Likelihood method under LG substitution model with a gamma distribution of rates between sites (eight categories, parameter alpha, estimated by the program) using PhyML 3.0 [21] with support values obtained by bootstrapping a 1000 replicates. For GenBank accession numbers of published NR sequences used in this study see **S1 file**.

## RESULTS

Genome databases of 33 species of Platyhelminths from Worm database were mined. The analyzed flatworm species include two species from class Rhabditophora, two species from class Monogenea, fifteen species from class Cestoda and fourteen species from class Trematoda (**Table 1**). For amino acid sequences of the DBD identified in these species see **S2 file**.

### 1. NRs in Platyhelminths

#### 1.1. Class: Monogenea

Most of the species in this class are ectoparasitic that live on the gills or skin of freshwater and marine fishes. Genome database of two species of Monogenea (*Protopolystoma xenopodis* [28] and *Gyrodactylus salaris* [27]) were analyzed.

##### 1.1.1. Order: Polyopisthocotylea

*Protopolystoma xenopodis* of Polystomatidae family was analyzed. *P. xenopodis* is a fluke of African clawed frogs its adult worms lives in the urinary bladder of the host. Twenty-three NRs were identified in *P. xenopodis*. Phylogenetic analysis shows they belong to five different NR subfamilies: seven members are from subfamily 1 (TR, E78, three HR96s and two divergent members), ten members are from subfamily 2 (HNF4, two RXRs, TR4, TLL, PNR, DSF, two Coup-TFs and NHR236), one member is from subfamily 4 (NR4A), two members are from subfamily 5 (FTZ-F1 and HR39) and three members belong to 2DBD-NRs (**Table 2, S1 Fig**).

**Table 2.**
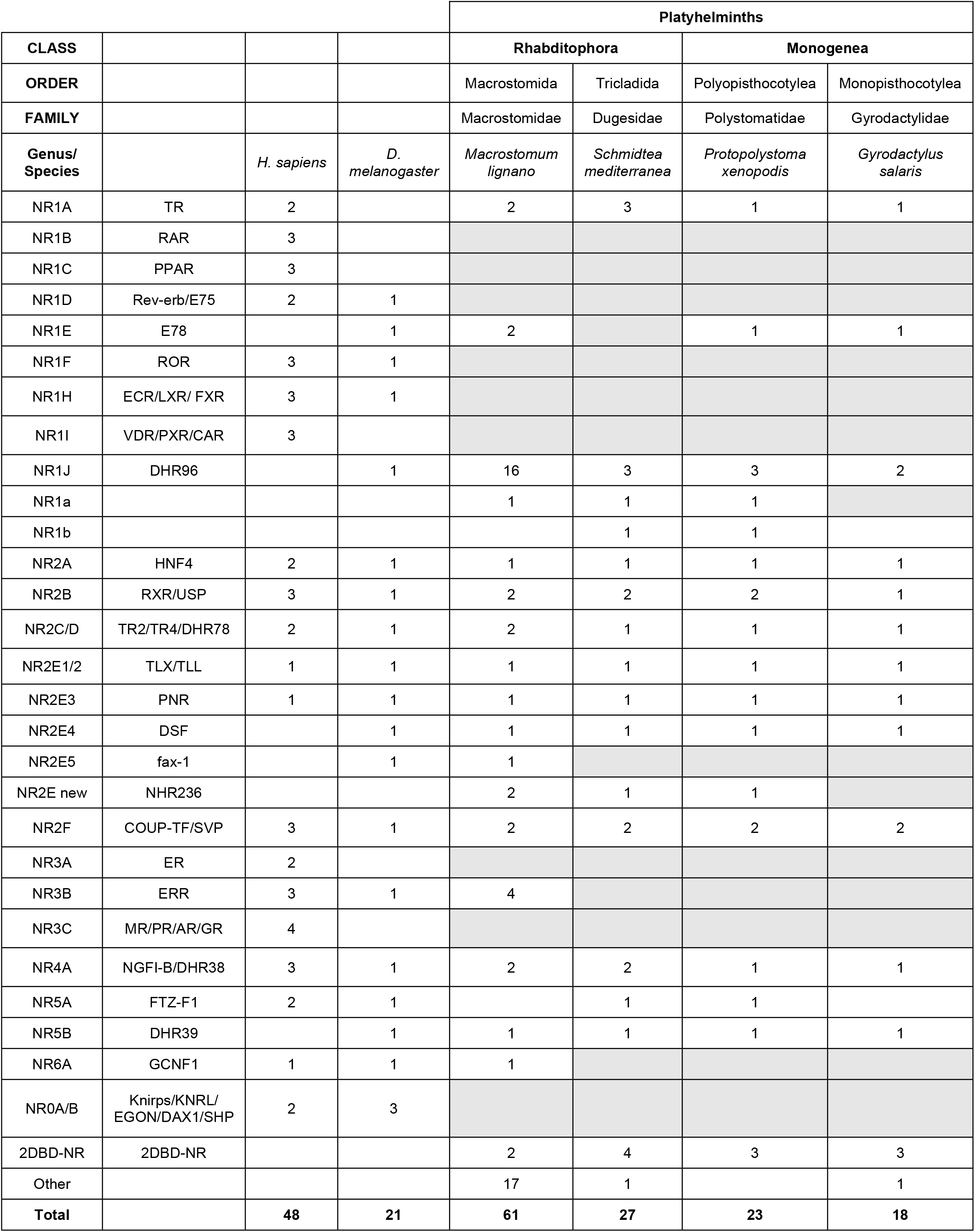
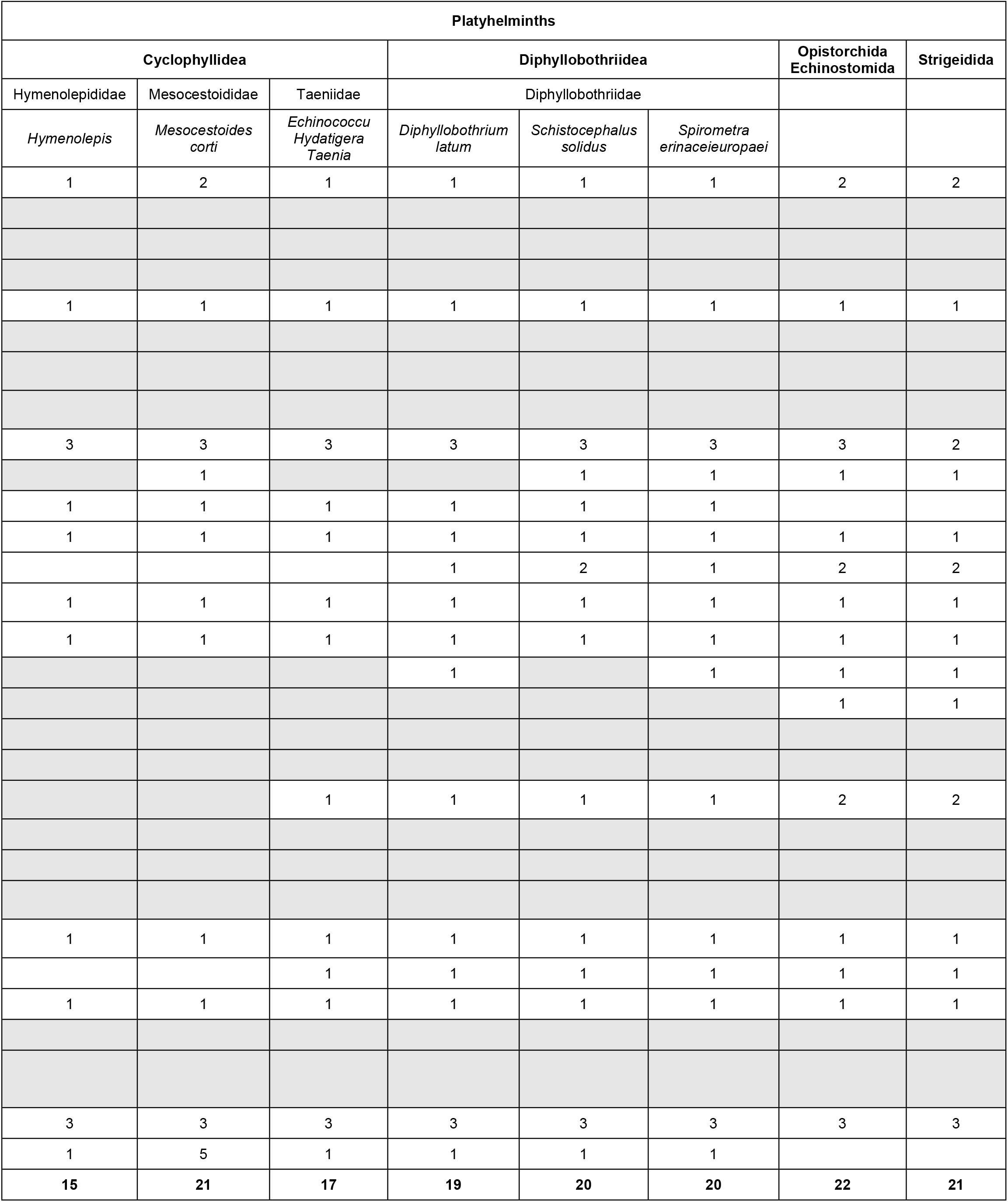
NRs in Platyhelminths.

One interesting result is that two divergent NRs from subfamily 1 (PxNR1a and PxNR1b) are present in *P. xenopodis*. Phylogenetic analysis shows that PxNR1a is an orthologue of *Schistosoma mansoni* NR1 (SmNR1) [44] (**S1 Fig**) and PxNR1b is an orthologue of the Cestode *Echinococcus granulosus* HR3 (EgHR3) [45]. Further analysis shows that the orthologue of PxNR1b is present in all analyzed tapeworms and also exists in the free-living Platyhelminth *Schmidtea mediterranea* (nhr10, [46]). Phylogenetic analysis of the DBD sequences shows that Platyhelminth NR1b group clustered together with *Drosophila* HR3/human ROR/Mollusca HR3 group but is outside of the HR3/ROR group (**S2 Fig**). To further demonstrate whether PxNR1b is an orthologue of ROR, DBD with LBD sequences or only LBD sequence of PxNR1b orthologue (EgHR3 and SmeNR1b) were analyzed. It turns out that the LBD sequence of PxNR1b is not available. Phylogenetic analysis results show that EgHR3 and SmeNR1b are clustered with the E75 group if both DBD and LBD sequences are analyzed, but they are clustered with E78 group if only LBD sequences are analyzed. These results suggest that PxNR1b orthologues are a group of divergent NRs that were present in a common Platyhelminth ancestor (**S3 – S4 Figs**).

Previously, we demonstrated that a novel NR2E member (orthologue of nematode *Caenorhabditis elegans* NHR236) was present in Cnidaria, Arthropoda, free-living Platyhelminths, Mollusca and Echinodermata [14]. In this study, an orthologue of NHR236 was identified in *P. xenopodis* (**Table 2, S1 Fig**). Phylogenetic analysis shows that NHR236 orthologues have a unique P-box sequence (CDGCRG) and they form a new NR subgroup in NR2E super group and its progenitor gene was present in a common ancestor of Porifera and Bilateral [14]. It is the first time that an orthologue of NHR236 has been shown to exist in parasitic Platyhelminths.

##### 1.1.2. Order Monopisthocotylea

*Gyrodactylus salaris* of Gyrodactylidae family was analyzed. *G. salaris* is a fish fluke which lives on the body surface of freshwater fish. *G. salaris* belongs to family Gyrodactylidae of order Monopisthocotylea. Eighteen NRs from four NR subfamilies were identified in *G. salaris*. They include four members from subfamily 1 (TR, E78 and two HR96s), eight members from subfamily 2 (HNF4, RXR, TR4, TLL, PNR, DSF and two Coup-TFs), one member from subfamily 4 (NR4A), one member from subfamily 5 (HR39), three 2DND-NRs and one divergent NR which has an unknown P-box sequence (EPCKV) (**Table 2, S5 Fig**). A comparison of the NR complement with that of *P. xenopodis*, shows there are three HR96s in *P. xenopodis* but only two in *G. salaris;* two RXRs are present in *P. xenopodis* but only one in *G. salaris*; two NR1 divergent members (PxNR1a and PxNR1b) and a NHR236 orthologue identified in *P. xenopodis* that is missing in *G. salaris*. A *G. salaris* divergent NR is not found in *P. xenopodis* (**Table 2**).

#### 1.2. Class Cestoda

##### 1.2.1. Order: Cyclophyllidea

###### 1.2.1.1. Hymenolepididae family

Three species in the genus *Hymenolepis* (*H. diminuta, H. microstoma* and *H. nana*) were analyzed. They are all rodent tapeworms and their intermediate hosts are arthropods. Fifteen NRs were identified in *H. diminuta* and *H. microstoma*. They include six members from subfamily 1 (TR, E78, three HR96s and NR1b), three NRs from subfamily 2 (HNF4, TR4 and TLL), one member from subfamily 4 (NR4A), one member from subfamily 5 (HR39), three 2DBD-NRs and one divergent NR. Fourteen NRs were identified in *H. nana*, a divergent NR identified in *H. diminuta* and *H. microstoma* is not present in *H. nana* **(Table 2, S2 file and S6 Fig)**.

###### 1.2.1.2. Mesocestoididae family

NR complement of *Mesocestoides corti* was mined and analyzed. *M. corti* is a mouse tapeworm, its intermediate hosts are birds and small mammals; its definitive host can be cats and dogs. Twenty-one NRs were identified in *M. corti*. There were eight members from subfamily 1 (two TRs, E78, three HR96s, NR1a and NR1b); three members from subfamily 2 (NHF4, TR4 and TLL); one member from subfamily 4 (NR4A); one member from subfamily 5 (HR39); three 2DND-NRs and five divergent NRs (**Table 2, S2 file, S7 Fig)**.

###### 1.2.1.3. Taeniidae family

NR complement in *Taeniidae* has been reported from the fox tapeworm *Echinococcus multilocularis* where seventeen NRs were identified [15]. In this study, we expanded the analysis to include three genera (*Echinococcus, Taenia* and *Hydatigera*), they are 1) *E. canadensis* (a canid tapeworm its intermediate hosts are cervid, camels and cattle and the definitive hosts are dogs and other canids). 2) *E. granulosus* (a dog tapeworm where intermediate hosts are humans and horses. The definitive hosts are dogs). 3) *E. multilocularis* (a fox tapeworm. Its intermediate hosts are rodents and the definitive hosts are foxes, coyotes, dogs and other canids). 4) *T. asiatica* (a human tapeworm whose intermediate hosts are pigs). 5) *T. multiceps* (a dog tapeworm whose intermediate hosts are sheep, goats and cattle). 6) *T. solium* (the pork tapeworm its intermediate hosts are pigs and the definitive hosts are humans). 7) *H. taeniaeformis* (a cat tapeworm its intermediate hosts are rodents).

Phylogenetic analysis result shows that all of above species share the same NR complement as *E. multilocularis* [15]; they exhibit six members from subfamily 1 (TR, E78, three HR96s and NR1b); four members from subfamily 2 (NHF4, TR4, TLL and Coup-TFI), one member from subfamily 4 (NR4A), two members from subfamily 5 (FTZ-F1and HR39), three 2DBD-NRs and a divergent NR. The divergent NR has a P-box of CDSCRA that is a homologue to SmRXR2 in the LBD [15] **(Table 2, S8, S9, S10 Figs)**.

##### 1.2.2. Order: Diphyllobothriidea

Three species from three genera of Diphyllobothriidae family were analyzed. They are 1) *Dibothriocephalus latus* (a human tapeworm its intermediate hosts are fish). 2) *Schitocephalus solidus* (a bird tapeworm). Its first intermediate hosts are cyclopoid and the second intermediate hosts are fish. Its definitive hosts are fish-eating water birds). 3) *Spirometra erinaceieuropaei* (a tapeworm its first intermediate hosts are cyclopoids and the second intermediate hosts are fish, reptiles, or other amphibians. Its definitive hosts are cats and dogs) (**Table 1**).

Nineteen NRs were identified in *D. latus* including six members from subfamily 1 (TR, E78, three HR96s and NR1b); five members from subfamily 2 (HNF4, RXR, TR4, TLL, PNR and Coup-TF); one member from subfamily 4 (NR4A); two members from subfamily 5 (FTZ-F1 and HR96), three 2DBD-NRs and one divergent NRs which is similar to Coup-TF with a different p-box sequence (**Table 2**).

Twenty NRs were identified in *S. solidus* including seven members from subfamily 1 (TR, E78, three HR96s, NR1a and NR1b); six members from subfamily 2 (HNF4, two RXRs, TR4, TLL and Coup-TF), one member from subfamily 4 (NR4A), two members from subfamily 5 members (FTZ-F1a and FTZ-F1b), three 2DBD-NRs and one divergent NRs which is an orthologue of *D. latus* divergent NR (**Table 2, S11 Fig)**.

Twenty NRs were identified in *S. erinaceieuropaei*. They include seven members from subfamily 1 (TR, E78, three HR96s, NR1a and NR1b); six members from subfamily 2 (HNF4, RXR, TR4, TLL, PNR and Coup-TF); one member from subfamily 4 (NR4A); two members from subfamily 5 members (FTZ-F1a and FTZ-F1b); three 2DBD-NRs and one divergent NRs which is an orthologue of *D. latus* and *S. solidus* divergent NR (**Table 2, S12 Fig**).

The difference of NR complement among *D. latus, S. solidus* and *S. erinaceieuropae* is that one RXR is identified in *D. latus* and *S. erinaceieuropae*, but two RXRs are found in *S. solidus;* One PNR is identified in *D. latus* and *S. erinaceieuropae* that is missing in *S. solidus*; NR1a and NR1b are in both *S. solidus* and *S. erinaceieuropae*, but NR1a is missing from *D. latus* (**Table 2**).

Comparison of the NR complement in Cestoda: Cyclophyllidea and Diphyllobothriidae orders, RXR and PNR are present in Diphyllobothriidae but they are missing in Cyclophyllidea.

#### 1.3. Class Trematoda

##### 1.3.1. Order: Strigeidida

NR complements in nine species of Schistosomatidae family were analyzed, they are 1) *Schistosoma mansoni* (a human blood fluke its intermediate hosts are freshwater snails). 2) *S. haematobium* (a human blood fluke its intermediate hosts are freshwater snails). 3) *S. japonicum* (a zoonotic blood fluke, its intermediate hosts are amphibious snails). 4) *S. curassoni* (a blood fluke, its intermediate hosts are freshwater snails and the definitive hosts are ruminants (cattle, goat, sheep). 5) *S. margrebowiei* (a blood fluke, its intermediate hosts are freshwater snails and the definitive hosts are antelope, buffalo and waterbuck). 6) *S. mattheei* (a blood fluke, its intermediate hosts are freshwater snails and the definitive hosts are bovids). 7) *S. rodhaini* (a blood fluke, its intermediate hosts are freshwater snails and the definitive host are rodents). 8) *S. bovis* (a blood fluke, its intermediate hosts are freshwater snails and the definitive hosts are ruminants (cattle, goats, sheep)). 9) *Trichobilharzia regent* (a neuropathogenic fluke, its intermediate host are freshwater snails and the definitive hosts are avian species).

NRs in parasitic trematodes were well described in *S. mansoni* [12-14, 16, 44, 47-62]. Twenty-one NRs were identified in *S. mansoni* including seven members from subfamily 1 (two TRs, E78, two DHR96s, NR1a (SmNR1 orthologue); nine members from subfamily 2 (HNF4, two RXRs, TR4, TLL, PNR, DSF and two Coup-TFs); one member from subfamily 4 (NR4A); two members from subfamily 5 (FTZ-F1 and HR39) and three 2DBD-NRs (**Table 2, S13, S14 Figs)**. Analysis of the Schistosomatidae species shows that all of them share the same NR complement as *S. mansoni*.

##### 1.3.2. Order: Opistorchida

NR complements in three species of Opistorchiidae family were analyzed. They are 1) *Clonorchis sinensis* (a human liver fluke, its first intermediate hosts are freshwater snails, the second intermediate hosts are freshwater fish and the definitive hosts are humans and other mammals). 2) *Opisthorchis viverrini* (a human liver fluke, its first intermediate hosts are freshwater snails, the second intermediate hosts are freshwater fish and the definitive hosts are humans and other mammals). 3) *O. felineus* (a cat liver fluke, its first intermediate hosts are freshwater snails, the second intermediate hosts are freshwater fish and the definitive hosts are humans and other mammals).

Twenty-two NRs are identified in each of above species (**Table 2, S15, S16 Figs)**. The NR complement in Opistorchida is same as that of Strigeidida, the only difference is that an additional HR96 is present Opistorchida.

##### 1.3.3. Order: Echinostomida

###### 1.3.3.1. Echinostomatidae family

NR complement of *Echinostoma caproni* were mined and analyzed. *E. caproni* is an intestinal fluke, its first intermediate hosts are freshwater snails and the second intermediate hosts are freshwater snails or frogs, humans can be definitive host. Twenty-two NRs were identified in *E. caproni*, its NR complement is the same as Opistorchida species (**Table 2, S17 Fig**).

###### 1.3.3.2. Fasciolidae family

NR complement of *Fasciola hepatica* were mined and analyzed. *F. hepatica* is a liver fluke, its intermediate hosts are freshwater snails and the definitive hosts are cattle, sheep and other mammals including humans. Twenty-two NRs were identified in *F. hepatica*, its NR complement is the same as *E. caproni* and Opistorchida species (**Table 2, S18 Fig**).

A comparison of Trematoda species, demonstrates that all of them share the same NR complement except that an HR96 is missing in Strigeidida.

#### 1.4. Class Rhabditophora

##### 1.4.1. Order: Macrostomida

*Macrostomum lignano* of Macrostomidae family was analyzed. *M. lignano* is a free-living, intertidal flatworm; it can regenerate most of its body parts. *M. lignano* is a new model organism for studying mechanisms of developmental and evolutionary biology [63]. Sixty-one NRs were identified in *M. lignano*. Phylogenetic analysis showed there are 44 typical NRs and 17 divergent NRs in this species.

The typical NRs in *M. lignano* include twenty-one members from subfamily 1 (two TRs, two E78s, sixteen HR96s and NR1a); thirteen members from subfamily 2 (HNF4, two RXRs, two TR4s, TLL, PNR, DSF, fax1, two NHR236s and two Coup-TFs); four members from subfamily 3 (four ERRs); two members from subfamily 4 (two NR4As); one member from subfamily 5 (HR96); one member from subfamily 6 (HR6) and two 2DBD-NRs (**Table 2, S19 Fig**).

An interesting result is that HR96 underwent multiple gene duplications in *M. lignano* and gave rise to sixteen HR96s (**Table 2, S19 Fig**). Another important result is that an orthologue of *Drosophila* fax-1 and four ERRs are identified in *M. lignano*, fax-1 and ERR are missing in all of the other analyzed Platyhelminth species. This is the first report that fax-1 and members from subfamily 3 are present in Platyhelminths (**Table 2, Fig. 1 and S19 Fig**).

**Figure 1.**
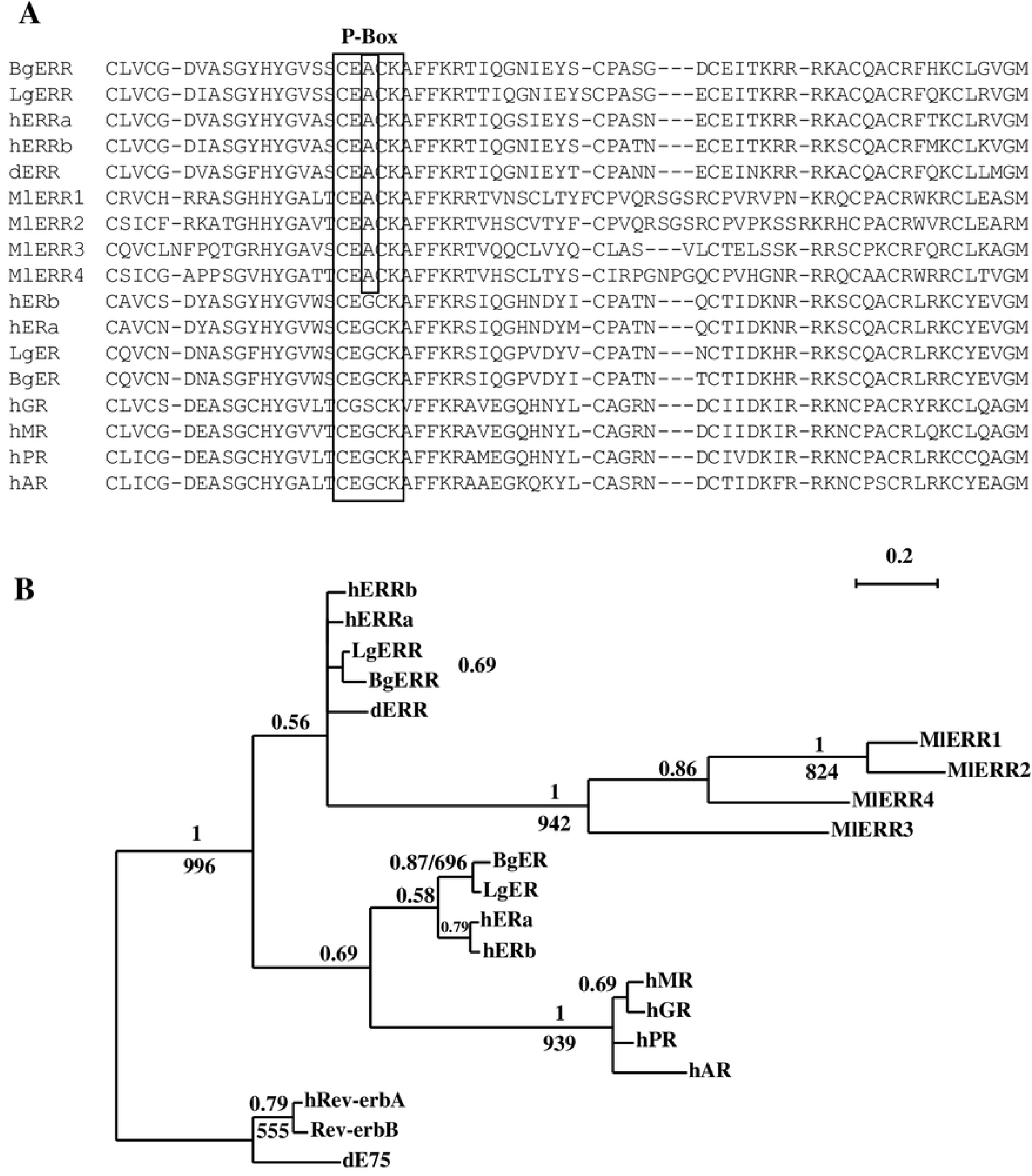
ERRs in Platyhelminths. **A)** Amino acid sequence alignment of DBD of MlERRs. **B)** Bayesian phylogenetic tree of MlERRs. The Bayesian tree is constructed with the deduced amino sequences of the DNA binding domain (DBD) with a mix amino acid replacement model + invgamma rates. The trees are started randomly with four simultaneous Markov chains running for 5 million generations. Bayesian posterior probabilities (BPPs) are calculated using a Markov chain Monte Carlo (MCMC) sampling approach implemented in MrBAYES v3.1.1, the PPs values are shown above each branch. Branches under the PPs 0.5 are shown as polytomies. The same data set is also tested by ML method using PHYML (v2.4.4) under LG substitution model (equilibrium frequencies model, proportion of invariable sites: estimated, number of substitution rate categories: 8, gamma shape parameter: estimated). Support values for the tree were obtained by bootstrapping a 1,000 replicates and bootstrap values above 500 and are indicated below each branch (or after MrBAYES BPPs separated by Slash). Star indicates the node obtained by Bayesian inference which is different from that obtained by ML method. Bg: *Biomphalaria glabrata*, d: *Drosophila melanogaster*, h: *Homo sapiens*, Lg: *Lottia gigantean*, Ml: *Macrostomum lignano*.

##### 1.4.2. Order Tricladida

*Schmidtea mediterranea* (Dugesidae family), a free-living, freshwater flatworm was analyzed. It can regenerate an entire organism and is an important model for stem cell research and regeneration [24]. Recently, twenty-three putative NRs were reported in *S. mediterranea* [46]. In this study, we re-analyze *S. mediterranea* genome database and twenty-seven NRs were identified, they include eight members from subfamily 1 (three TRs, three HR96s, NR1a and NR1b); ten members from subfamily 2 (HNF4, two RXRs, TR4, TLL, PNR, DSF, NHR236 and two Coup-TFs); two members from subfamily 4 (NR4As); two members from subfamily 5 (FTZ-F1 and HR96), four 2DBD-NRs and a divergent NR (**Table 2, S20 Fig)**.

### 2. A novel zinc finger motif (CHC2) exists in DBD of parasitic Platyhelminth NRs

DBD is the most conserved region in NRs, it contains two C4-type zinc finger motifs. In each motif, four cysteine residues chelate one Zn^2+^ ion. The first zinc finger (CI) contains a sequence element, the P-box [64, 65] which is responsible for binding the target gene, and the second zinc finger (CII) contains a sequence element the D-box which is responsible for dimerization [64].

Previously, we isolated a partial cDNA of a trematode *Schistosoma mansoni* HR96b (SmNR96b) [12], unlike other NRs, SmNR96b contains a long amino acid sequence in DBD with two introns located in this region. DBD of SmHR96b has a histidine residue replaced by a second cysteine residue in the D-Box in CII and it forms a novel zinc finger motif (CHC2). Recently, CHC2 zinc finger motif has been demonstrated to be present in HR96b of Cestoda and other Trematoda species [66, 67]. In this study, we determined that all SmHR96b orthologues in parasitic Platyhelminths contain this novel CHC2 zinc finger motif (**Fig. 2**). Thus, CHC2 type zinc finger motif represents a novel NR CII which has diverged in ancient HR96b in a common ancestor of parasitic Platyhelminths. The function of this novel CHC2 type motif in NRs is unknown. Recent study shows that SmHR96b (named Vitellogenic Factor 1 in [66, 67]) is essential for female sexual development.

**Figure 2.**
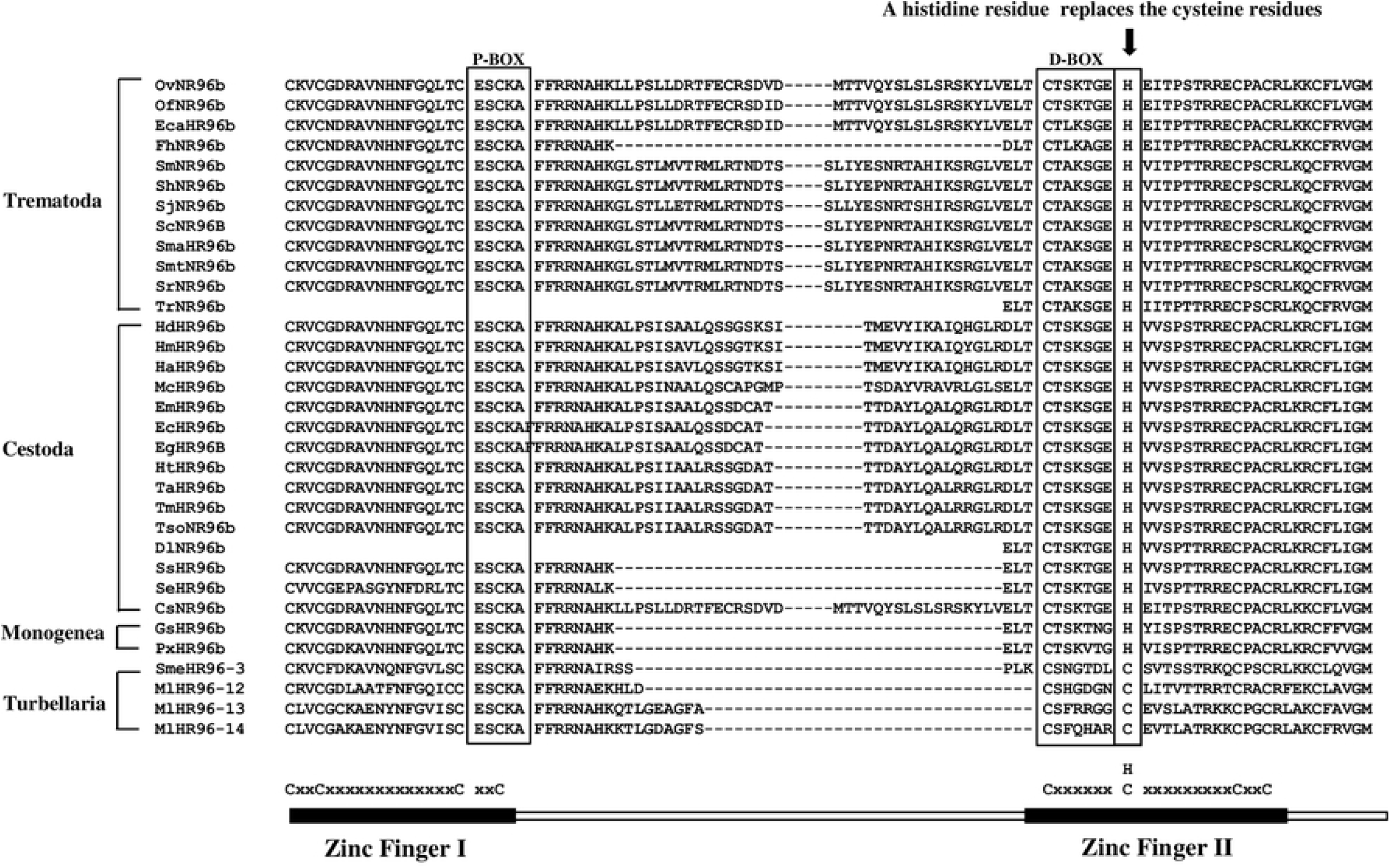
A novel zinc finger motif (CHC2) in DBD of parasitic Platyhelminth HR96b. Amino acid sequence alignment of DBD of the Platyhelminth HR96b shows a novel NR zinc finger motif (CHC2) is present in parasitic Platyhelminth NRs. A histidine residue replaces the second cysteine residues in the D-Box of zinc finger II forming a novel CHC2 motif. See Fig 1 legend for methods for phylogenetic tree construction. Cs: *Clonorchis sinensis*, Dl: *Dibothriocephalus latus*, Ec: *Echinococcus canadensis*, Eca: *Echinostoma caproni*, Eg: *Echinococcus granulosus*, Em: *Echinococcus multilocularis*, Fh: *Fasciola hepatica*, Gs: *Gyrodactylus salaris*, Hd: *Hymenolepis diminuta*, Hn: *Hymenolepis nana*, Ht: *Hydatigera taeniaeformis*, Lg: *Lottia gigantean*, Mc: *Mesocestoides corti*, Ml: *Macrostomum lignano*, Of: *Opisthorchis felineus*, Ov: *Opisthorchis viverrini*, Px: *Protopolystoma xenopodis*, Sb: *Schistosoma bovis*, Sc: *Schistosoma curassoni*, Se: *Spirometra erinaceieuropaei*, Sh: *Schistosoma haematobium*, Sj: *Schistosoma japonicum*, Sm: *Schistosoma mansoni*, Sma: *Schistosoma margrebowiei*, Smt: *Schistosoma mattheei*, Sme: *Schmidtea mediterranea*, Sr: *Schistosoma rodhaini*, Ss: *Schitocephalus solidus*, Ta: *Taenia asiatica*, Tm: *Taenia multiceps*, Ts: *Taenia saginata*, Tr: *Trichobilharzia regent*, Tso: *Taenia solium*.

### 3. Divergent NRs

Divergent NR refers to a NR which has a typical P-box sequence in DBD but it does not fall into any ‘typical’ NR groups, for example Platyhelminths NR1a and NR1b with a typical P-box of ‘CEGCKGFFRR’ belonging to NR subfamily 1 but they do not fit into any groups within the NR1 subfamily. It also refers to a NR which has an atypical P-box sequence in DBD and does not fall into the present NR nomenclature [68].

Divergent NRs exist in various lineages of Platyhelminths. One divergent NR with a typical P-box (CEGCKGFFKR) which is same as that of RXR/TR4/NR4A is found in Rhabditophora *S. mediterranea* and in Cestoda *Mesocestoides corti*. Most of the Platyhelminth divergent NRs possess an ‘atypical’ P-box sequence. For example, a NR with a P-box of ‘CEPCKVFFKR’ is identified in Monogenea *G. salaris*, a NR with a P-box of ‘CEACKAFFQQ’ is found in all analyzed species of Cestoda Hymenolepis family, a NR which has a P-box of ‘CDSCRAFFEM’ exists in Cestoda Taenia family, a NR with a P-box of ‘CEACKSFFKR’ is found in Cestoda Diphyllobothriidea order and seventeen NRs with various ‘atypical’ P-box sequences were identified in Rhabditophora *M. lignano* (**S21Fig**).

A phylogenetic tree of the divergent NRs including those of Platyhelminths and Mollusca was constructed (**S21Fig**). Phylogenetic analysis shows no orthologues exist among Rhabditophora *M. lignano*, Cestoda and Mollusca. The result suggests that these divergent NRs diverged and duplicated independently in different animal lineages.

### 4. Evolution of NR gene in Platyhelminths

The number of NRs varies in different Platyhelminth species, e.g. 27-62 members are found in Rhabditophora, 18-23 members are found in Monogenea, 15-20 members are found in Cestoda and 21-22 members are found in Trematoda. This variety is a result of gene duplication/amplification and gene loss in different Platyhelminth lineages.

#### 4.1. Subfamily 1

##### 4.1.1. E78

E78 was present as a common ancestor of Platyhelminths. It was retained in Rhabditophora *M. lignano*, Monogenea, Cestoda and Trematoda, but it was missing in Rhabditophora *S. mediterranea*. E78 underwent a duplication in *M. lignano* and gave a birth of two E78s (MlE78a and MlE78b).

##### 4.1.2. TR

One TR is identified in Monogenea; two are identified in Rhabditophora *M. lignano* and three are identified in *S. mediterranea*; and two are identified in Cestoda and Trematoda, respectively. Our previous study showed that two TR homologues are present in Platyhelminths [12-14, 61]. Phylogenetic analysis suggested that Platyhelminth TR gene duplicated after the split of the trematodes and the turbellarians [61], thus the paralogue of the trematode TR was not present in a turbellarian and in turn the paralogue of turbellarian TR was not present in trematode species.

Phylogenetic analysis in this study shows that all TRs of parasitic Platyhelminths are clustered in a group, two *M. lignano* TRs are clustered in a group and three *S. mediterranea* TRs are clustered in another group. This result suggests that TRs duplicated independently in *M. lignano, S. mediterranea* and parasitic Platyhelminths. In parasitic Platyhelminth TR groups, trematode TRa (orthologues of *Schistosoma* TRa, SmTRa) are clustered together with those of Monogenea and Cestoda, but trematode TRb (orthologues of Schistosoma TRb, SmTRb) group only contains trematode TRs. This result suggested that one TR was present in a common ancestor of Platyhelminths and trematode TRa was an ancient TR gene. Phylogenetic analysis shows that the two Cestode *M. corti* TRs are clustered together in Cestoda TR group, this result suggests that TRs in Cestoda and Trematoda underwent duplication independently (**S22 Fig)**. Since TR duplicated independently in different Platyhelminths lineage, Platyhelminths TRb/TRc are paralogous genes.

##### 4.1.3. HR96

Sixteen HR96s are identified in Rhabditophora *M. lignano* and two are found in *S. mediterranea*; three HR96s are identified in Monogenea, Cestoda and Trematoda, respectively.

Phylogenetic analysis shows that Platyhelminthes HR96s are clustered in four different groups: HR96a, HR96b, HR96c and HR96d. HR96a, HR96b and HR96c groups contain only Platyhelminth HR96s, but HR96d group contains both Platyhelminth and Mollusca members. The result suggests that Platyhelminth HR96d is an ancient NR gene (**Fig. 3**). HR96d group only contains Platyhelminth *M. lignano* HR96s, but each group of HR96a, HR96b and HR96c contains NR96s from all species of the four classes of Platyhelminths, this result suggests that there were four HR96s in a common ancestor of Platyhelminths and the ancient HR96 gene (HR96d) was lost in *Schmidtea mediterranea*, Monogenea, Cestoda and Trematoda (**Fig. 3**). HR96 genes were amplified in *M. lignano* and each of the four HR96 genes gave birth to a total of sixteen NR96 genes.

**Fig. 3.**
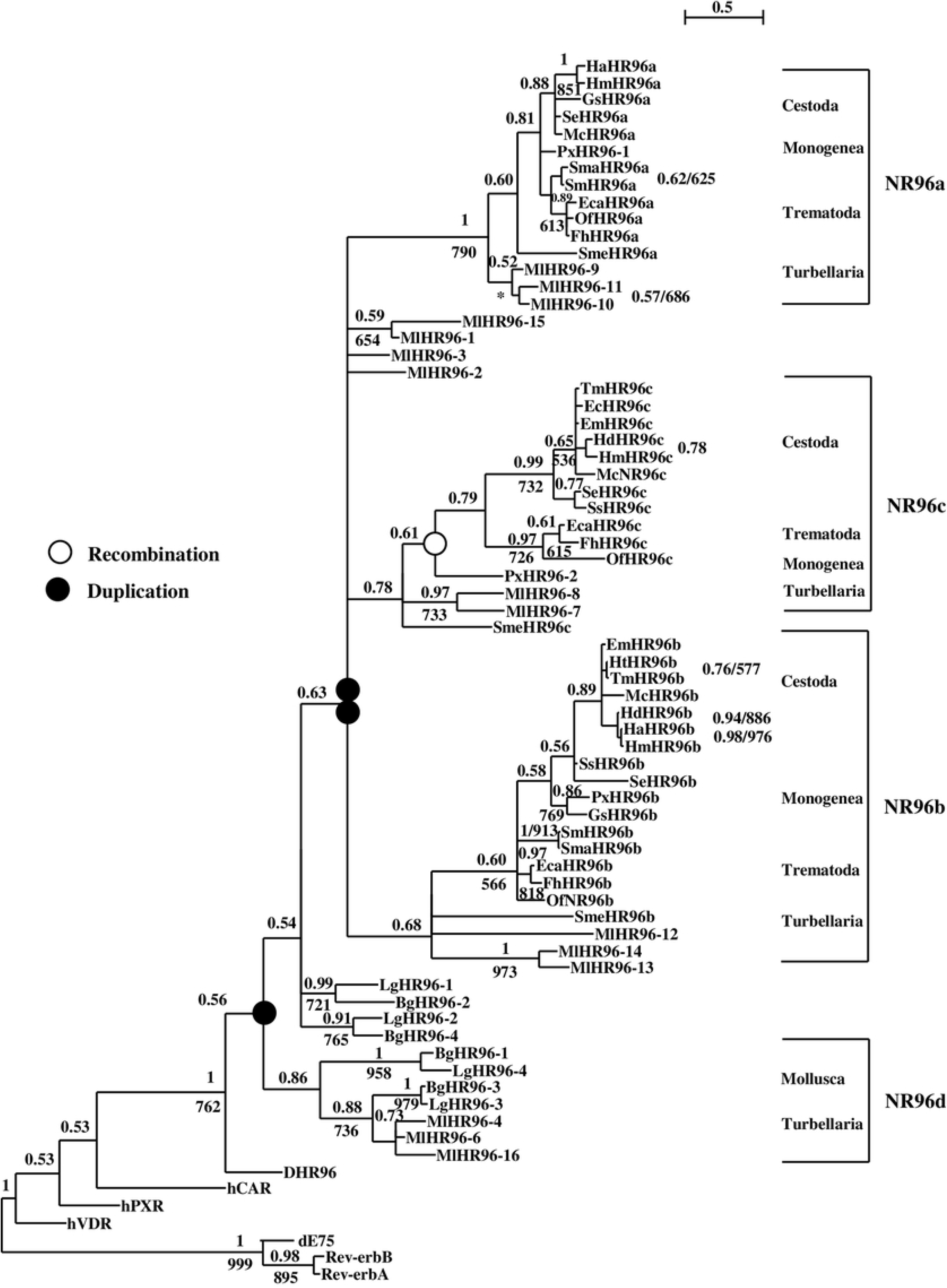
Bayesian phylogenetic tree of Platyhelminth HR96s. Methods for construction of phylogenetic trees see figure 1 legend. Bg: *Biomphalaria glabrata*, Cs: *Clonorchis sinensis*, d: *Drosophila melanogaster*, Dl: *Dibothriocephalus latus*, Ec: *Echinococcus Canadensis*, Eca: *Echinostoma caproni*, Eg: *Echinococcus granulosus*, Em: *Echinococcus multilocularis*, Fh: *Fasciola hepatica*, Gs: *Gyrodactylus salaris*, h: *Homo sapiens*, Hd: *Hymenolepis diminuta*, Hn: *Hymenolepis nana*, Ht: *Hydatigera taeniaeformis*, Lg: *Lottia gigantean*, Mc: *Mesocestoides corti*, Ml: *Macrostomum lignano*, Of: *Opisthorchis felineus*, Ov: *Opisthorchis viverrini*, Px: *Protopolystoma xenopodis*, Sb: *Schistosoma bovis*, Sc: *Schistosoma curassoni*, Se: *Spirometra erinaceieuropaei*, Sh: *Schistosoma haematobium*, Sj: *Schistosoma japonicum*, Sm: *Schistosoma mansoni*, Sma: *Schistosoma margrebowiei*, Smt: *Schistosoma mattheei*, Sme: *Schmidtea mediterranea*, Sr: *Schistosoma rodhaini*, Ss: *Schitocephalus solidus*, Ta: *Taenia asiatica*, Tm: *Taenia multiceps*, Ts: *Taenia saginata*, Tr: *Trichobilharzia regent*, Tso: *Taenia solium*.

##### 4.1.4. Divergent NR1a

Parasitic Platyhelminth NR1a gene is an orthologue of *S. mansoni* NR1 (SmNR1) [44]. It is retained in Monogenea *P. xenopodis*, Cestoda *S. solidus* and *S. erinaceieuropaei* and Trematoda. NR1a was missing in Monogenea *G. salaris*, Cestoda *D. latum*, and Hymenolepididae and Taeniidae families. One divergent NR1a is also present in Rhabditophora *M. lignano* and *S. mediterranea*. Phylogenetic analysis cannot determine if they are orthologues of parasitic Platyhelminth NR1a.

##### 4.1.5. Divergent NR1b

Platyhelminths NR1b gene is an orthologue of EgHR3 [45]. It is retained in Rhabditophora *S. mediterranea*, Monogenea *P. xenopodis* and all analyzed tapeworms, it was missing in Rhabditophora *M. lignano*, Monogenea *G. salaris* and Trematoda.

#### 4.2. Subfamily 2

##### 4.2.1. HNF4

HNF4 retained in all analyzed Platyhelminths species.

##### 4.2.2. RXR

Two RXRs were identified in Rhabditophora, Monogenea, Cestoda and Trematoda, respectively. Phylogenetic analysis shows that parasitic flatworm RXRs are clustered in two different groups: RXR1 group (*Schistosoma* RXR1 orthologues) and RXR2 group (*Schistosoma* RXR2 orthologues). RXR2 group is clustered with RXRs of free-living flatworms, *Drosophila*, Mollusca and human, this suggests that parasitic Platyhelminth RXR2 is an orthologue of free-living flatworms, *Drosophila*, Mollusca and human RXRs. RXR1 group contains only parasitic Platyhelminth RXRs, this suggests that parasitic Platyhelminth RXR2 duplicated after the split of their common ancestor with free-living Platyhelminths. In free-living flatworms, two *M. lignano* RXRs are clustered in a group while two *S. mediterranea* RXRs are scattered in the phylogenetic tree, this result suggests that RXR duplicated independently in free-living flatworms *M. lignano* and *S. mediterranea*. The intron position in DBD of RXRs further supports this result (**S23 Fig**). RXR gene is lost in some of parasitic Platyhelminths after ancient RXR duplicated in a common ancestor of parasitic Platyhelminths: RXR1 was lost in Monogenea *G. salaris* and Cestoda *D. latum*; RXR2 was lost in Cestoda *S. erinaceieuropaei*; both RXR1 and RXR2 were lost in all the analyzed species of Cestoda Cyclophyllidea order.

##### 4.2.3. TR4

TR4 is present in all analyzed Platyhelminth species except two members are present in Rhabditophora *M. lignano* (MlTR4a and MLTR4b). This suggests that TR4 gene underwent duplication in *M. lignano*.

##### 4.2.4. TLL

TLL is a conserved NR member, it is present in all analyzed Platyhelminth species.

##### 4.2.5. PNR

PNR was lost in Cestoda Cyclophyllidea order and Cestoda *S. solidus*. It is retained in all other analyzed Platyhelminth species.

##### 4.2.6. DSF

DSF is absent in Cestoda.

##### 4.2.7. fax-1

One fax1 was identified in Rhabditophora *M. lignano*, but was missing in all other analyzed Platyhelminths.

##### 4.2.8. NHR236

Two NHR236s were present in Rhabditophora *M. lignano*, one is present in Rhabditophora *S. mediterranea* and Monogenea *P. xenopodis*. It was lost in Monogenea *G. salaris*, Cestoda and Trematoda.

##### 4.2.9. Coup-TF

One Coup-TF was identified in Cestoda; two are identified in Rhabditophora, Monogenea and Trematoda, respectively. Phylogenetic analysis shows that parasitic flatworms Coup-TFs are clustered in two different groups: Coup-TFI group (*Schistosoma* Coup-TFI orthologues) and Coup-TFII group (*Schistosoma* Coup-TFII orthologues). Coup-TFI group contains members from all three parasitic classes and it is clustered with that of free-living flatworms, *Drosophila*, Mollusca and humans. This result suggests that parasitic Platyhelminth Coup-TFI is an ancient Coup-TF gene and it is the orthologue of free-living flatworms, *Drosophila*, Mollusca and human Coup-TFs. Parasitic Platyhelminth Coup-TFII group contains Monogenea and Trematoda members, this suggests that Coup-TFI duplicated and gave rise to Coup-TFII in a common ancestor of parasitic Platyhelminths. In free-living flatworms, two *M. lignano* Coup-TFs and two *S. mediterranea* Coup-TFs are scattered in the phylogenetic tree, it suggests *M. lignano* and *S. mediterranea* Coup-TFs duplicated independently; analysis of the intron position in DBD of Coup-TFs further supports this result (**S24 Fig**). Coup-TFI was missing in Cestoda Hymenolepididae family and in Mesocestoididae (*M. corti*); Coup-TFII was missing in all analyzed species of Cestoda.

#### 4.3. Subfamily 3

Four ERRs were identified in Rhabditophora *M. lignano*. Phylogenetic analysis shows that all four *M. lignano* ERRs form single group clustered with *Drosophila*, Mollusca and human ERRs. The result demonstrates that *M. lignano* ERRs duplicated after a split of *Drosophila*, Mollusca, Platyhelminths and humans (**Fig. 3B**). ERR is missing in Rhabditophora *S. mediterranea*, and in all analyzed parasitic Platyhelminths.

#### 4.4. Subfamily 4

Two NR4As were identified in Rhabditophora *M. lignano* and *S. mediterranea*, respectively; one is found in all analyzed parasitic flatworms. Phylogenetic analysis shows that MlNR4Aa, SmeNR4Aa and parasitic flatworm NR4A are clustered with *Drosophila*, Mollusca and human NR4A; it suggests that MlNR4Aa, SmeNR4Aa and parasitic flatworm NR4A are orthologues of *Drosophila*, Mollusca and human NR4A. MlNR4Ab, SmeNR4Ab are scattered in the phylogenetic tree suggesting that they underwent a duplication after a split of *M. lignano* and *S. mediterranea* (**S25 Fig**).

#### 4.5. Subfamily 5

##### 4.5.1. FTZ-F1

FTZ-F1 is missing in Rhabditophora *M. lignano*, Monogenea *G. salaris*, Cestoda Hymenolepididae and Mesocestoididae families. It is retained in all the other analyzed Platyhelminths (**S26 Fig**).

##### 4.5.2. HR39

HR39 is retained in all analyzed Platyhelminths (**S26 Fig**).

##### 4.6. Subfamily 6

NR6 is present in Rhabditophora *M. lignano* but lost in Rhabditophora *S. mediterranea* and all analyzed parasitic Platyhelminths.

##### 4.7. 2DBD-NR

Two 2DBD-NRs were identified in Rhabditophora, *M. lignano* and four were identified in *S. mediterranea*; three 2DBD-NRs were found in parasitic Platyhelminths including Monogenea, Cestoda and Trematoda, respectively.

A phylogenetic tree of 2DBD-NRs was constructed with the amino acid sequence of the second DBD, because the second DBD is more conserved than the first DBD. Phylogenetic analysis shows that parasitic Platyhelminth 2DBD-NRs clustered in three groups: 2DBDa (*Schistosoma mansoni* 2DBDα orthologues), 2DBDb (*S. mansoni* 2DBDβ orthologues) and 2DBDg (*S. mansoni* 2DBDγ orthologues) groups. Parasitic Platyhelminth 2DBD-NRa and 2DBD-NRb groups contain members of three classes of parasitic flatworms, but 2DBD-NRg group contains 2DBD-NRs of all the four classes of Platyhelminths and also members of the Mollusca. This result suggests that Parasitic Platyhelminth 2DBD-NRg is an ancient gene, and 2DBD-NRa and 2DBD-NRb were formed by a second round of duplication. In free living Platyhelminths, both Rhabditophora *M. lignano* 2DBD-NRs are clustered in 2DBD-NRg group, but the four *S. mediterranea* 2DBD-NRs are clustered with different parasitic helminth 2DBD-NR groups. This result suggests that *M. lignano* 2DBD-NR gene underwent duplication after a split of *S. mediterranea* and parasitic Platyhelminths. For the four *S. mediterranea* 2DBD-NRs, one is clustered in parasitic Platyhelminth 2DBD-NRg group, one is clustered in 2DBD-NRb group and two are clustered with 2DBD-NRa group. Since the two *S. mediterranea* (2DBD-NRa1 and 2DBD-NRa2) in 2DBDa group form a polytomy, it suggests that 2DBD-NRa underwent another round of duplication and formed two 2DBD-NRs as a common ancestor of *S. mediterranea* and parasitic Platyhelminths and then one 2DBD-NRa was lost in a common ancestor of parasitic Platyhelminths (**S27Fig**).

The above results suggests that there were at least twenty-four ancient NRs in a common ancestor of Platyhelminths (**Fig. 4**), there were eight members from subfamily 1 (TR, E78, DHR96a, DHR96b, DHR96c, DHR96d, NR1a and NR1b); ten members from subfamily 2 (HNF4, RXR2, TR4, TLL, PNR, DSF, dax1, NHR236, Coup-TFI and Coup-TFII); one member from subfamily 3 (ERR); One member from subfamily 4 (NR4A); two members from subfamily 5 (FTZ-F1 and HR39); one member from subfamily 6 (GRF); one member from 2DBD-NRs (2DBD-NRg). Phylogenetic analysis shows that NR gene gain/loss occurred in different Platyhelminth lineages (**Fig. 4**).

**Figure 4.**
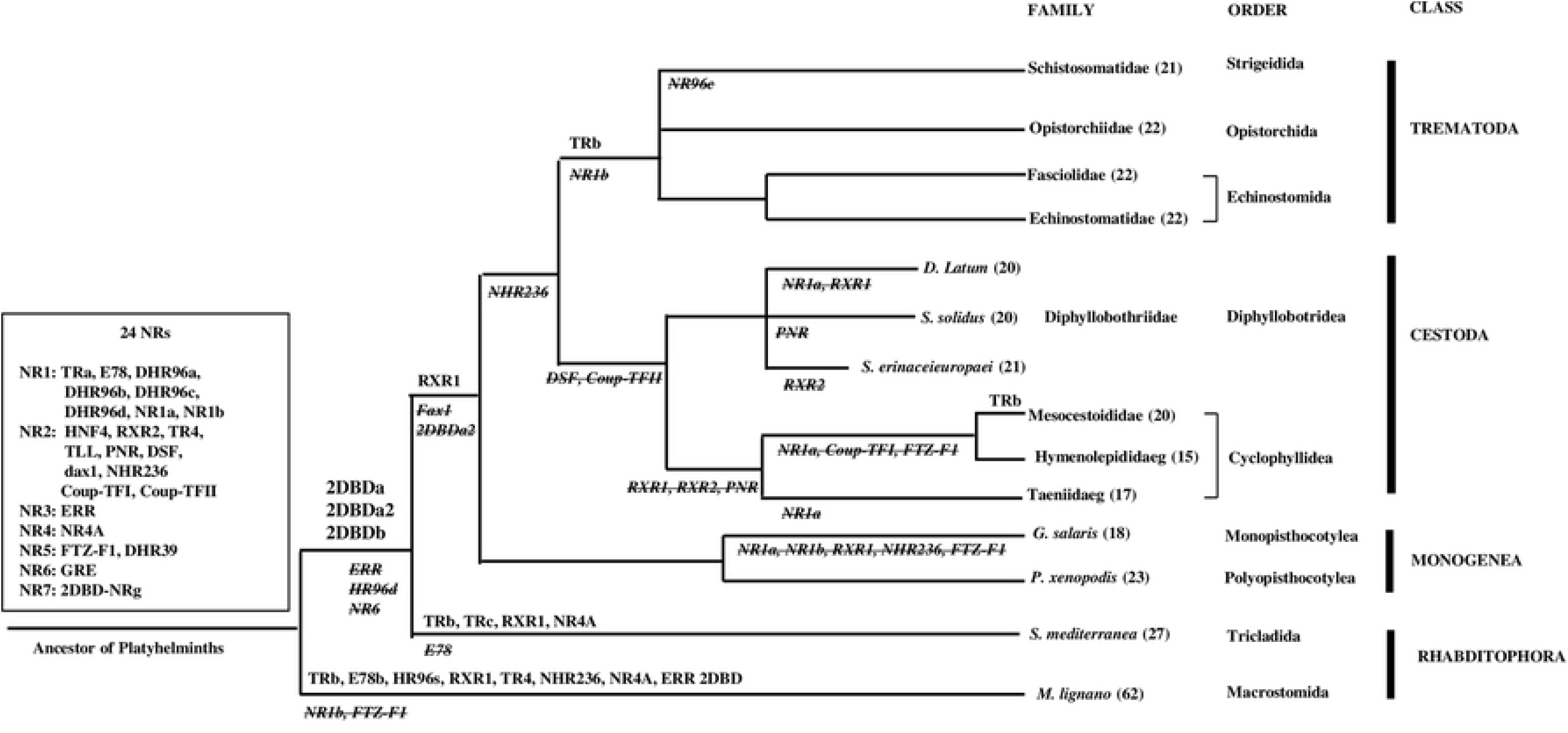
Summary of NR gene gain and loss in the evolution of Platyhelminths. A scheme represents the phylogeny of Platyhelminths. The twenty-four ancient NR genes in a common ancestor of Platyhelminths are indicated in square box on the left of the figure. NR gene gained by gene duplication is shown above the branch of different flatworm lineages and the gene lost is shown under the branches (italic and strikethrough). The number behind flatworm species/families in parentheses indicates the number of NRs identified.

## DISCUSSION

In this study, we performed a phylogenetic analysis of the evolution of NRs in Platyhelminths. The results show that NRs in Platyhelminths have orthologues in Deuterostomes, arthropods or both, and the NRs in Platyhelminths diverged into two different evolutionary lineages: 1) Gene duplication and lost; 2) NR gene amplification and divergence.

NRs in Rhabditophora *S. mediterranea* and parasitic Platyhelminths are followed in the first evolutionary lineage: gene duplication and lost, the events occurred in different flatworm lineages. For example, fax-1 was lost in a common ancestor of *S. mediterranea* and parasitic Platyhelminths, NHR236 was lost in endoparasites (Cestoda and Trematoda), and DSF was lost in Cestoda. In parasitic Platyhelminths, extensive NR gene loss occurred in Cestoda Hymenolepididae and Taeniidae families, e. g. RXR, Coup-TF and FTZ-F1 were retained in most species of Platyhelminths, but they were missing in the cestode families Hymenolepididae and Taeniidae.

Comparison of NR complement in different Platyhelminth families, NR complement is most conserved in Trematoda. All species of Trematoda share the same NR complement except that a NHR96 member was lost in Strigeidida (*Schistosoma* and *Trichobilharzia*). NR complement in Monogenea and in Cestoda exhibit more differences among different families, in Monogenea, there are eighteen NRs in Gyrodactylidae (*G. salaris*), but twenty-three NRs are present in Polyopisthocotylea (*P. xenopodis*). In Cestoda Cyclophyllidea, there are fifteen NRs in Hymenolepididae family, seventeen in Taeniidae family and twenty in Mesocestoididae family. NR gene duplication occurred in different flatworm lineages. For example, 2DBD-NR underwent a second round of duplication in a common ancestor of *S. mediterranea* and parasitic Platyhelminths, RXR gene duplicated separately in *S. mediterranea* and a common ancestor of parasitic Platyhelminths, and TR duplicated independently in Rhabditophora (*S. mediterranea*), Cestoda and Trematoda.

NRs in Rhabditophora *M. lignano* follow another evolutionary lineage: Gene amplification and divergence. In *M. lignano*, sixty-one NRs are identified including sixteen HR96s and seventeen divergent members. This result suggests that NRs in *M. lignano* follows an evolutionary lineage with a feature of multiple gene duplication (amplification) and gene divergence. Multiple NR gene duplication was also found in other animals, for example, supplementary NRs (SupNRs) in nematodes and NR1H in Chordate *Cephalochordate amphioxus* [69, 70].

NRs in subfamily 3 and 6 were considered lost in Platyhelminths. In this study, we identified four ERRs and a NR6A in Rhabditophora *M. lignano*. This is the first known occurrence of NRs in subfamily 3 and 6 in Platyhelminths, but it is still not clear whether NRs from subfamily 3 and 6 are present in other Rhabditophora since the genome data is unavailable. This study also shows that divergent NRs are present in different flatworm lineages suggesting that novel NRs were acquired in different flatworms to adapt to the different living environments.

Previously, we isolated a partial cDNA of *S. mansoni* HR96b (SmNR96b) [12], this member has a CHC2 zinc finger motif in the second zinc finger of DBD. In this study, we show that all parasitic Platyhelminth SmHR96b orthologues contain this novel motif. Whether the function of this new type of motif in NRs may change the DNA binding properties awaits further study.

## Acknowledgments

We acknowledge Kevin L. Howe, Bruce J. Bolt, Myriam Shafie, Paul Kersey, and Matthew Berriman for access to **WormBase ParaSite – a comprehensive resource for helminth genomics**.

## Abbreviation

2DBD-NR: nuclear receptor with two DBDs
AR: androgen receptor
CAR: constitutive androstane receptor
COUP-TF: chicken ovalbumin upstream promoter transcription factor
DBD: DNA binding domain
DSF: *Drosophila* dissatisfaction gene
E75: *Drosophila* ecdysone-induced protein 75
E78: *Drosophila* ecdysone-induced protein 78
EAR: V-erbA-related gene
EGON: embryonic gonad protein
ER: estrogen receptor
ERR: estrogen related receptor
FTZ-F1: fushi tarazu-factor 1
FXR: farnesoid X receptor
GCNF: germ cell nuclear factor
GR: glucuronide receptor
HNF-4: hepatocyte nuclear factor 4
HR96: *Drosophila* hormone receptor 96
LBD: ligand binding domain
LRH-1: liver receptor homologue 1
LXR: liver X receptor
MR: mineralocorticoid receptor
NGFI-B: nerve growth factor-induced B
NHR236: nematode nuclear receptor 236
NR: Nuclear receptor
Nurr1: nuclear receptor related 1
NOR1: neuron-derived orphan receptor 1
PNR: photoreceptor-specific nuclear receptor
PR: progesterone receptor
PXR: pregnane X receptor
PPAR: peroxisome proliferators-activated receptor
Rev-erb: nuclear receptor-related protein coded on the opposite strand of the thyroid hormone receptor gene
RAR: retinoic acid receptor
ROR: retinoid-related orphan receptor
RXR: retinoid X receptor
SF1: Steroidogenic factor 1
SR: steroid receptor
SVP: seven up; Supplementary nuclear receptors
SupNRs: supplementary nuclear receptors
TLL: *Drosophila* tailless gene
TLX: homologue of the *Drosophila* tailless gene
TR: thyroid hormone receptor
TR2/4: testicular receptor 2/4
USP: ultraspiracle
VDR: vitamin D receptor

## Supplemental FIGURE LEGENDS

**S1 Fig. Bayesian phylogenetic tree of *Protopolystoma xenopodis* NRs**

The Bayesian tree was constructed with the deduced amino sequences of the DNA binding domain (DBD) with a mix amino acid replacement model + invgamma rates. The trees were started randomly with four simultaneous Markov chains running for 5 million generations. Bayesian posterior probabilities (BPPs) are calculated using a Markov chain Monte Carlo (MCMC) sampling approach implemented in MrBAYES v3.1.1, the PPs values are shown above each branch. Branches under the PPs 0.5 are shown as polytomies. The same data set is also tested by ML method using PHYML (v2.4.4) under LG substitution model (equilibrium frequencies model, proportion of invariable sites: estimated, number of substitution rate categories: 8, gamma shape parameter: estimated). Support values for the tree are obtained by bootstrapping a 1,000 replicates and bootstrap values above 500 and are indicated below each branch (or after MrBAYES BPPs separated by Slash). Star indicates the node obtained by Bayesian inference which is different from that obtained by ML method. Bg: *Biomphalaria glabrata*, Cg: *Crassostrea gigas*, d: *Drosophila melanogaster*, h: *Homo sapiens*, Lg: *Lottia gigantean*, Px: *Protopolystoma xenopodis*, Sm: *Schistosoma mansoni*. Red highlighted NRs show *P. xenopodis* NRs.

**S2 Fig. Bayesian phylogenetic tree of NR1 DBD sequences**

Methods for construction of phylogenetic trees see S1 fig legend. Bg: *Biomphalaria glabrata*, d: *Drosophila melanogaster*, Dl: *Dibothriocephalus latus*, Eca: *Echinostoma caproni*, Eg: *Echinococcus granulosus*, Fh: *Fasciola hepatica*, h: *Homo sapiens*, Hd: *Hymenolepis diminuta*, Ht: *Hydatigera taeniaeformis*, Lg: *Lottia gigantean*, Mc: *Mesocestoides corti*, Ml: *Macrostomum lignano*, Ov: *Opisthorchis viverrini*, Px: *Protopolystoma xenopodis*, Se: *Spirometra erinaceieuropaei*, Sm: *Schistosoma mansoni*, Sme: *Schmidtea mediterranea*, Ss: *Schitocephalus solidus*, Tm: *Taenia multiceps*, Tr: *Trichobilharzia regent*. Red highlighted NRs show Platyhelminths NRs.

**S3 Fig. Bayesian phylogenetic tree of NR1 DBD with LBD**

Methods for construction of phylogenetic trees see S1 fig legend. Bg: *Biomphalaria glabrata*, d: *Drosophila melanogaster*, Eg: *Echinococcus granulosus*, h: *Homo sapiens*, Lg: *Lottia gigantean*, Sme: *Schmidtea mediterranea*. Red highlighted NRs show Platyhelminths NRs.

**S4 Fig. Bayesian phylogenetic tree of NR1 LBD**

Methods for construction of phylogenetic trees see S1 fig legend. Bg: *Biomphalaria glabrata*, d: *Drosophila melanogaster*, Eg: *Echinococcus granulosus*, h: *Homo sapiens*, Lg: *Lottia gigantean*, Sme: *Schmidtea mediterranea*. Red highlighted NRs show Platyhelminths NRs.

**S5 Fig. Bayesian phylogenetic tree *Gyrodactylus salaris* NRs**

Methods for construction of phylogenetic trees see S1 fig legend. Bg: *Biomphalaria glabrata*, Cg: *Crassostrea gigas*, d: *Drosophila melanogaster*, Gs: *Gyrodactylus salaris*, h: *Homo sapiens*, Lg: *Lottia gigantean*, Px: *Protopolystoma xenopodis*, Sm: *Schistosoma mansoni*. Red highlighted NRs show *G. salaris* NRs.

**S6 Fig. Bayesian phylogenetic tree of *Hymenolepis microstoma* NRs**

The phylogenetic tree of *H. microstoma* NRs represents all analyzed *Hymenolepis* species because of the highly conserved DBD sequences in these species. Methods for construction of phylogenetic trees see S1 fig legend. Bg: *Biomphalaria glabrata*, Cg: *Crassostrea gigas*, d: *Drosophila melanogaster*, h: *Homo sapiens*, Hm: *Hymenolepis microstoma*, Lg: *Lottia gigantean*, Px: *Protopolystoma xenopodis*, Sm: *Schistosoma mansoni*. Red highlighted NRs show *H. microstoma* NRs.

**S7 Fig. Bayesian phylogenetic tree of *Mesocestoides corti* NRs**

Methods for construction of phylogenetic trees see S1 fig legend. Bg: *Biomphalaria glabrata*, Cg: *Crassostrea gigas*, d: *Drosophila melanogaster*, h: *Homo sapiens*, Lg: *Lottia gigantean*, Mc: *Mesocestoides corti*, Px: *Protopolystoma xenopodis*, Sm: *Schistosoma mansoni*. Red highlighted NRs show *M. corti* NRs.

**S8 Fig. Bayesian phylogenetic tree of *Echinococcus multilocularis* NRs**

Methods for construction of phylogenetic trees see S1 fig legend. Bg: *Biomphalaria glabrata*, Cg: *Crassostrea gigas*, d: *Drosophila melanogaster*, h: *Homo sapiens*, Lg: *Lottia gigantean, Em: Echinococcus multilocularis*, Px: *Protopolystoma xenopodis*, Sm: *Schistosoma mansoni*. Red highlighted NRs show *E. multilocularis* NRs.

**S9 Fig. Bayesian phylogenetic tree of *Hydatigera taeniaeformis* NRs**

Methods for construction of phylogenetic trees see S1 fig legend. Bg: *Biomphalaria glabrata*, Cg: *Crassostrea gigas*, d: *Drosophila melanogaster*, h: *Homo sapiens*, Ht: *Hydatigera taeniaeformis*, Lg: *Lottia gigantean*, Px: *Protopolystoma xenopodis*, Sm: *Schistosoma mansoni*. Red highlighted NRs show *H. taeniaeformis* NRs.

**S10 Fig. Bayesian phylogenetic tree of *Taenia saginata* NRs**

Methods for construction of phylogenetic trees see S1 fig legend. Bg: *Biomphalaria glabrata*, Cg: *Crassostrea gigas*, d: *Drosophila melanogaster*, h: *Homo sapiens*, Lg: *Lottia gigantean*, Px: *Protopolystoma xenopodis*, Sm: *Schistosoma mansoni*, Ts: *Taenia saginata*. Red highlighted NRs show *T. saginata* NRs.

**S11 Fig. Bayesian phylogenetic tree of *Schitocephalus solidus* NRs**

Methods for construction of phylogenetic trees see S1 fig legend. Bg: *Biomphalaria glabrata*, Cg: *Crassostrea gigas*, d: *Drosophila melanogaster*, h: *Homo sapiens*, Ss: *Schitocephalus solidus*, Lg: *Lottia gigantean*, Px: *Protopolystoma xenopodis*, Sm: *Schistosoma mansoni*. Red highlighted NRs show *T. saginata* NRs.

**S12 Fig. Bayesian phylogenetic tree of *Spirometra erinaceieuropaei* NRs**

Methods for construction of phylogenetic trees see S1 fig legend. Bg: *Biomphalaria glabrata*, Cg: *Crassostrea gigas*, d: *Drosophila melanogaster*, h: *Homo sapiens*, Lg: *Lottia gigantean*, Px: *Protopolystoma xenopodis*, Se: *Spirometra erinaceieuropaei*, Sm: *Schistosoma mansoni*. DBD of E78 sequence is partial, not used for construction of the tree. Red highlighted NRs show *S. erinaceieuropaei* NRs.

**S13 Fig. Bayesian phylogenetic tree of *Schistosoma haematobium* NRs**

The phylogenetic tree of *S haematobium* NRs represents all analyzed *Schistosoma* species because of the highly conserved DBD sequences in these species. Methods for construction of phylogenetic trees see S1 fig legend. Bg: *Biomphalaria glabrata*, Cg: *Crassostrea gigas*, d: *Drosophila melanogaster*, h: *Homo sapiens*, Lg: *Lottia gigantean*, Px: *Protopolystoma xenopodis*, Sh: *Schistosoma haematobium*, Sm: *Schistosoma mansoni*. Red highlighted NRs show *S. haematobium* NRs.

**S14 Fig. Bayesian phylogenetic tree of *Trichobilharzia regent* NRs**

Methods for construction of phylogenetic trees see S1 fig legend. Bg: *Biomphalaria glabrata*, Cg: *Crassostrea gigas*, d: *Drosophila melanogaster*, h: *Homo sapiens*, Lg: *Lottia gigantean*, Px: *Protopolystoma xenopodis*, Sm: *Schistosoma mansoni*, Tr: *Trichobilharzia regent*. Red highlighted NRs show *T. regent* NRs.

**S15 Fig. Bayesian phylogenetic tree of *Clonorchis sinensis* NRs**

Methods for construction of phylogenetic trees see S1 fig legend. Bg: *Biomphalaria glabrata*, Cg: *Crassostrea gigas*, Cs: *Clonorchis sinensis*, d: *Drosophila melanogaster*, h: *Homo sapiens*, Lg: *Lottia gigantean*, Px: *Protopolystoma xenopodis*, Sm: *Schistosoma mansoni*. Red highlighted NRs show *C. sinensis* NRs.

**S16 Fig. Bayesian phylogenetic tree of *Opisthorchis viverrini* NRs**

The phylogenetic tree of *O viverrini* NRs represents *Opisthorchis viverrini* and *O. felineus* because of the highly conserved DBD sequences in these two species. Methods for construction of phylogenetic trees see S1 fig legend. Bg: *Biomphalaria glabrata*, Cg: *Crassostrea gigas*, d: *Drosophila melanogaster*, h: *Homo sapiens*, Lg: *Lottia gigantean*, Ov: *Opisthorchis viverrini*, Px: *Protopolystoma xenopodis*, Sm: *Schistosoma mansoni*. Red highlighted NRs show *O. viverrini* NRs.

**S17 Fig. Bayesian phylogenetic tree of *Echinostoma caproni* NRs**

Methods for construction of phylogenetic trees see S1 fig legend. Bg: *Biomphalaria glabrata*, Cg: *Crassostrea gigas*, d: *Drosophila melanogaster*, Eca: *Echinostoma caproni*, h: *Homo sapiens*, Lg: *Lottia gigantean*, Px: *Protopolystoma xenopodis*, Sm: *Schistosoma mansoni*. EcaCoup-TFII sequence is partial and not used for construction of the tree. Red highlighted NRs show *E. caproni* NRs.

**S18 Fig. Bayesian phylogenetic tree of *Fasciola hepatica* NRs**

Methods for construction of phylogenetic trees see S1 fig legend. Bg: *Biomphalaria glabrata*, Cg: *Crassostrea gigas*, d: *Drosophila melanogaster*, Fh: *Fasciola hepatica*, h: *Homo sapiens*, Lg: *Lottia gigantean*, Px: *Protopolystoma xenopodis*, Sm: *Schistosoma mansoni*. Red highlighted NRs show *F. hepatica* NRs.

**S19 Fig. Bayesian phylogenetic tree of *Macrostomum lignano* NRs**

Methods for construction of phylogenetic trees see S1 fig legend. Bg: *Biomphalaria glabrata*, Cg: *Crassostrea gigas*, d: *Drosophila melanogaster*, h: *Homo sapiens*, Lg: *Lottia gigantean*, Ml: *Macrostomum lignano*, Px: *Protopolystoma xenopodis*, Sm: *Schistosoma mansoni*. Red highlighted NRs show *M. lignano* NRs.

**S20 Fig. Bayesian phylogenetic tree of *Schmidtea mediterranea* NRs**

Methods for construction of phylogenetic trees see S1 fig legend. Bg: *Biomphalaria glabrata*, Cg: *Crassostrea gigas*, d: *Drosophila melanogaster*, h: *Homo sapiens*, Lg: *Lottia gigantean*, Px: *Protopolystoma xenopodis*, Sm: *Schistosoma mansoni*, Sme: *Schmidtea mediterranea*. Red highlighted NRs show *S. mediterranea* NRs.

**S21 Fig. Bayesian phylogenetic tree of Platyhelminth divergent NRs**

Methods for construction of phylogenetic trees see S1 fig legend. Bg: *Biomphalaria glabrata*, d: *Drosophila melanogaster*, Dl: *Dibothriocephalus latus*, Ec: *Echinococcus Canadensis*, Eg: *Echinococcus granulosus*, Em: *Echinococcus multilocularis*, Gs: *Gyrodactylus salaris*, h: *Homo sapiens*, Hd: *Hymenolepis diminuta*, Hm: of *H. microstoma*, Ht: *Hydatigera taeniaeformis*, Mc: *Mesocestoides corti*, Ml: *Macrostomum lignano*, Sme: *Schmidtea mediterranea*, Ta: *Taenia asiatica*, Tm: *Taenia multiceps*, Ts: *Taenia saginata*, Tr: *Trichobilharzia regent*, Tso: *Taenia solium*.

**S22 Fig. Bayesian phylogenetic tree of Platyhelminth TRs**

Methods for construction of phylogenetic trees see S1 fig legend. Bg: *Biomphalaria glabrata*, Cs: *Clonorchis sinensis*, d: *Drosophila melanogaster*, Dl: *Dibothriocephalus latus*, Ec: *Echinococcus Canadensis*, Eca: *Echinostoma caproni*, Eg: *Echinococcus granulosus*, Em: *Echinococcus multilocularis*, Fh: *Fasciola hepatica*, Gs: *Gyrodactylus salaris*, h: *Homo sapiens*, Hd: *Hymenolepis diminuta*, Hm: of *H. microstoma*, Hn: *Hymenolepis nana*, Ht: *Hydatigera taeniaeformis*, Lg: *Lottia gigantean*, Mc: *Mesocestoides corti*, Ml: *Macrostomum lignano*, Of: *Opisthorchis felineus*, Ov: *Opisthorchis viverrini*, Px: *Protopolystoma xenopodis*, Sb: *Schistosoma bovis*, Sc: *Schistosoma curassoni*, Se: *Spirometra erinaceieuropaei*, Sh: *Schistosoma haematobium*, Sj: *Schistosoma japonicum*, Sm: *Schistosoma mansoni*, Sma: *Schistosoma margrebowiei*, Smt: *Schistosoma mattheei*, Sme: *Schmidtea mediterranea*, Sr: *Schistosoma rodhaini*, Ss: *Schitocephalus solidus*, Ta: *Taenia asiatica*, Tm: *Taenia multiceps*, Ts: *Taenia saginata*, Tr: *Trichobilharzia regent*, Tso: *Taenia solium*.

**S23 Fig. Bayesian phylogenetic tree of Platyhelminth RXR**

A) Bayesian phylogenetic tree of Platyhelminth RXR. Methods for construction of phylogenetic trees see S1 fig legend. Bg: *Biomphalaria glabrata*, Cs: *Clonorchis sinensis*, d: *Drosophila melanogaster*, Dl: *Dibothriocephalus latus*, Ec: *Echinococcus Canadensis*, Eca: *Echinostoma caproni*, Eg: *Echinococcus granulosus*, Em: *Echinococcus multilocularis*, Fh: *Fasciola hepatica*, Gs: *Gyrodactylus salaris*, h: *Homo sapiens*, Lg: *Lottia gigantean*, Ml: *Macrostomum lignano*, Of: *Opisthorchis felineus*, Ov: *Opisthorchis viverrini*, Px: *Protopolystoma xenopodis*, Sb: *Schistosoma bovis*, Sc: *Schistosoma curassoni*, Se: *Spirometra erinaceieuropaei*, Sh: *Schistosoma haematobium*, Sj: *Schistosoma japonicum*, Sm: *Schistosoma mansoni*, Sma: *Schistosoma margrebowiei*, Smt: *Schistosoma mattheei*, Sme: *Schmidtea mediterranea*, Sr: *Schistosoma rodhaini*, Ss: *Schitocephalus solidus*, Tr: *Trichobilharzia regent*. B) Sequence alignment shows the intron position (red color >) in DBD of RXRs.

**S24 Fig. Bayesian phylogenetic tree of Platyhelminth Coup-TF**

A) Bayesian phylogenetic tree of Platyhelminth Coup-TF. Methods for construction of phylogenetic trees see S1 fig legend. Bg: *Biomphalaria glabrata*, Cs: *Clonorchis sinensis*, d: *Drosophila melanogaster*, Dl: *Dibothriocephalus latus*, Ec: *Echinococcus Canadensis*, Eca: *Echinostoma caproni*, Eg: *Echinococcus granulosus*, Em: *Echinococcus multilocularis*, Fh: *Fasciola hepatica*, Gs: *Gyrodactylus salaris*, h: *Homo sapiens*, Lg: *Lottia gigantean*, Ml: *Macrostomum lignano*, Of: *Opisthorchis felineus*, Ov: *Opisthorchis viverrini*, Px: *Protopolystoma xenopodis*, Sb: *Schistosoma bovis*, Sc: *Schistosoma curassoni*, Se: *Spirometra erinaceieuropaei*, Sh: *Schistosoma haematobium*, Sj: *Schistosoma japonicum*, Sm: *Schistosoma mansoni*, Sma: *Schistosoma margrebowiei*, Smt: *Schistosoma mattheei*, Sme: *Schmidtea mediterranea*, Sr: *Schistosoma rodhaini*, Ss: *Schitocephalus solidus*, Ta: *Taenia asiatica*, Tm: *Taenia multiceps*, Ts: *Taenia saginata*, Tr: *Trichobilharzia regent*, Tso: *Taenia solium*. B) Sequence alignment shows the intron position (red color >) in DBD of Coup-TFs.

**S25 Fig. Bayesian phylogenetic tree of Platyhelminth NR4A**

Methods for construction of phylogenetic trees see S1 fig legend. Bg: *Biomphalaria glabrata*, d: *Drosophila melanogaster*, Gs: *Gyrodactylus salaris*, h: *Homo sapiens*, Hd: *Hymenolepis diminuta*, Lg: *Lottia gigantean*, Mc: *Mesocestoides corti*, Ml: *Macrostomum lignano*, Sm: *Schistosoma mansoni*, Sme: *Schmidtea mediterranea*.

**S26 Fig. Bayesian phylogenetic tree of subfamily 5**

Methods for construction of phylogenetic trees see S1 fig legend. Bg: *Biomphalaria glabrata*, d: *Drosophila melanogaster*, Dl: *Dibothriocephalus latus*, Ec: *Echinococcus Canadensis*, Eca: *Echinostoma caproni*, Em: *Echinococcus multilocularis*, Fh: *Fasciola hepatica*, Gs: *Gyrodactylus salaris*, h: *Homo sapiens*, Hn: *Hymenolepis nana*, Lg: *Lottia gigantean*, Mc: *Mesocestoides corti*, Ml: *Macrostomum lignano*, Of: *Opisthorchis felineus*, Ov: *Opisthorchis viverrini*, Px: *Protopolystoma xenopodis*, Se: *Spirometra erinaceieuropaei*, Sj: *Schistosoma japonicum*, Sm: *Schistosoma mansoni*, Smt: *Schistosoma mattheei*, Sme: *Schmidtea mediterranea*, Tr: *Trichobilharzia regent*, Tso: *Taenia solium*.

**S27 Fig. Bayesian phylogenetic tree of Platyhelminth 2DBD**

Methods for construction of phylogenetic trees see S1 fig legend. Bg: *Biomphalaria glabrata*, Cs: *Clonorchis sinensis*, d: *Drosophila melanogaster*, Dl: *Dibothriocephalus latus*, Em: *Echinococcus multilocularis*, Fh: *Fasciola hepatica*, Gs: *Gyrodactylus salaris*, h: *Homo sapiens*, Hd: *Hymenolepis diminuta*, Hm: *H. microstoma*, Lg: *Lottia gigantean*, Mc: *Mesocestoides corti*, Ml: *Macrostomum lignano*, Px: *Protopolystoma xenopodis*, Se: *Spirometra erinaceieuropaei*, Sm: *Schistosoma mansoni*, Sme: *Schmidtea mediterranea*, Ss: *Schitocephalus solidus*, Tr: *Trichobilharzia regent*.

